# *In planta* transcriptomics reveals conflicts between pattern-triggered immunity and the AlgU sigma factor regulon

**DOI:** 10.1101/2022.01.03.474781

**Authors:** Haibi Wang, Amelia Lovelace, Amy Smith, Brian H. Kvitko

**Affiliations:** Department of Plant Pathology, University of Georgia, Athens GA; The Sainsbury Laboratory, Norwich Research Park, Norwich, NR4 7UH, UK; The Plant Center, University of Georgia, Athens GA

**Author notes:** Corresponding author: B.H.Kvitko. HW contributed to data analysis and writing, AL contributed to sequencing data processing, AS contributed to RNA sample preparation and sequencing, BK oversaw the project. Funding: National Science Foundation Division of Integrative Organismal Systems grant number 1844861 to B. H. Kvitko.

**Keywords:** *Arabidopsis thaliana*, *Pseudomonas syringae*, plant immunity, PTI, *algU*, T3E, motility, transcriptome

## Abstract

In previous work, we determined the transcriptomic impacts of flg22 pre-induced Pattern Triggered Immunity (PTI) in *Arabidopsis thaliana* on the pathogen *Pseudomonas syringae* pv. *tomato* DC3000 (*Pto*). During PTI exposure we observed expression patterns in *Pto* reminiscent of those previously observed in a *Pto algU* mutant. AlgU is a conserved extracytoplasmic function sigma factor which has been observed to regulate over 950 genes in *Pto in vitro*. We sought to identify the AlgU regulon *in planta*.and which PTI-regulated genes overlapped with AlgU-regulated genes. In this study, we analyzed transcriptomic data from RNA-sequencing to identify the AlgU *in planta* regulon and its relationship with PTI. Our results showed that approximately 224 genes are induced by AlgU, while another 154 genes are downregulated by AlgU in *Arabidopsis* during early infection. Both stress response and virulence-associated genes were induced by AlgU, while the flagellar motility genes are downregulated by AlgU. Under the pre-induced PTI condition, more than half of these AlgU-regulated genes have lost induction/suppression in contrast to naïve plants, and almost all function groups regulated by AlgU were affected by PTI.

---

The plant cell produces surface receptors that allow it to recognize Pathogen Associated Molecular Patterns (PAMPs) and trigger Pattern Triggered Immunity (PTI). In the model organism *Arabidopsis thaliana* Col-0, PTI induction is associated with a series of responses including a rapid ROS (reactive oxygen species) burst, induction of defensive hormone biosynthesis pathways, changes in apoplast and cell wall composition, and increased expression of pathogen-related receptors (DeFalco and Zipfel 2021; Jwa and Hwang 2017; Ngou et al. 2021; Luna et al. 2011). The flagellin-derived peptide flg22 is a synthetic PAMP commonly used to activate PTI in laboratory settings. Upon recognizing flg22, the *Arabidopsis* FLS2 receptor initiates PTI responses, reducing proliferation of the model pathogen *Pseudomonas syringae* pv. *tomato* DC3000 (*Pto*) (Zipfel et al. 2004). An activated PTI response also restricts Type III effector delivery by *Pto* and reduces expression of the *P. syringae* virulence regulon (Crabill et al. 2010; Lovelace et al. 2018; O’Malley and Anderson 2021; Nobori et al. 2018).

Upon transitioning into the leaf apoplast *Pto* has the capacity to perceive and respond to niche-specific cues to reprogram gene expression and express plant virulence factors. Extracytoplasmic function (ECF) sigma factors are a common tool used by bacteria to achieve such reprogramming. One ECF sigma factor, AlgU (RpoE), is known to regulate hundreds of genes involved in metabolism, motility, stress tolerance, and virulence in *Pto* (Markel et al. 2016). Deletion of the *algU* gene or the *algUmucAB* operon, which includes the anti-sigma factors of AlgU, reduces *Pto* growth in tomato and *Arabidopsis* seedlings (Markel et al. 2016; Ishiga et al. 2018).

Previous studies have identified *Pto* genes with altered regulation during exposure to pre-induced PTI in *Arabidopsis* including motility related genes, osmotic stress response genes, and alginate synthesis genes (Lovelace et al. 2018; Nobori et al. 2018). Although these genes overlap with the known *in vitro* AlgU regulon, it is not known to what degree the AlgU *in planta* regulon (AlgU-regulated genes in the plant-niche) resembles the *in vitro* AlgU regulon (AlgU-regulated genes in growth media), or how PTI modifies the AlgU *in planta* regulon. In this study, we identified the *Pto* AlgU regulon during early interactions with the *Arabidopsis* adult leaf apoplast, and in addition show that pre-induced PTI intervenes against the induction or suppression of more than half of these AlgU-regulated genes.

## Results

### Transcriptomic differences between *Pto* WT and *ΔalgU* during early *Arabidopsis* infection reveals the AlgU *in planta* regulon

In order to identify the AlgU *in planta* regulon, we infiltrated *Arabidopsis* with either wild type (WT) or *ΔalgUmucAB* (*ΔalgU*) *Pto*, and collected total RNA samples at 5 hpi (hours post inoculation) for RNA-seq using previously established techniques (Lovelace et al. 2018). This time point was chosen to be consistent with said study. After plant host and bacterial rRNA depletion, samples were sequenced by RNA-Seq. The resulting reads were cleaned and mapped to the *Pto* reference genomes and differentially-expressed genes (DEGs) were identified for both strains. By using a cutoff at 1.3-fold change in expression and false discovery rate less than 5%, we found that 224 genes were upregulated, while 154 genes were down regulated in the WT background compared to the *ΔalgU* background (Fig. 1A and Table S1 and S2). To confirm our observations, we validated the expression change patterns of genes belong to function groups of interest including alginate production (*algD*), osmotic stress response (*opuCA*), motility (*fliD*), and virulence (*avrPto1*) by Reverse Transcription-quantitative PCR (RT-qPCR) (Fig. S1). It is worth noting that since some of these genes are encoded in polycistronic operons, the number of different transcripts regulated by AlgU is smaller than the number of the identified genes.

**Figure 1.**
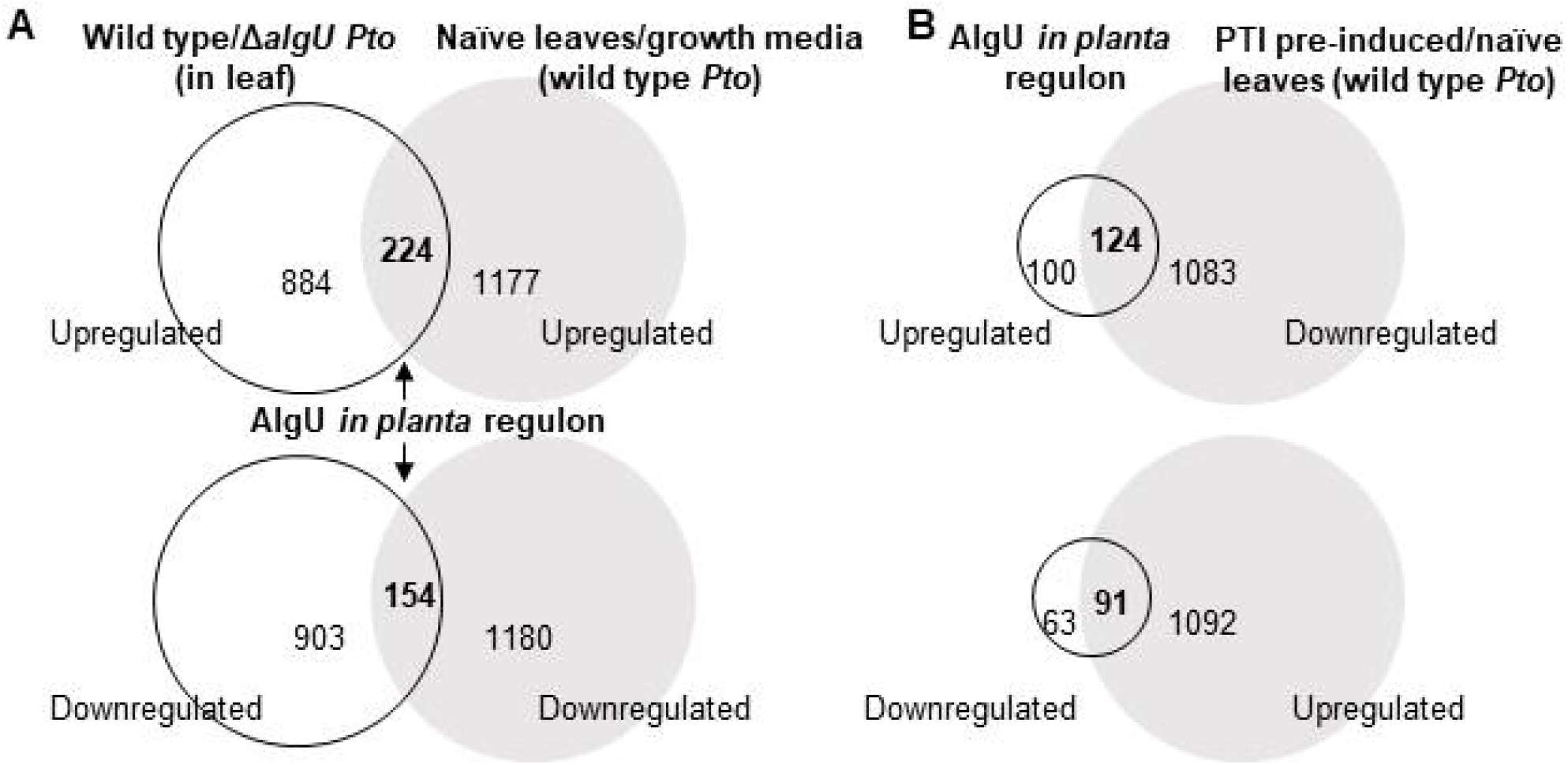
Venn diagram showing numbers of genes differentially-regulated under different conditions. **A**. AlgU *in planta* regulon genes is present in both left(circle): upregulated (top) or downregulated (bottom) in wild type compared to *ΔalgU Pto*, and right(grey): upregulated (top) or downregulated (bottom) in naïve leaves compared to King’s B growth media. **B**. Proportion of genes upregulated (top) or downregulated (bottom) by AlgU in plant (circle) overlaps with genes that are expressed more in mock-treated leaves (naïve) compared to flg22-treated (PTI pre-induced) leaves (grey).

For the AlgU upregulated *in planta* regulon, we were able to identify genes involved in osmotic stress response, oxidative stress response, alginate synthesis and export, cell shape and division pathways, DNA repair pathways, Type II Secretion System structural genes, and macromolecule metabolism and energy production (including lipid metabolism, amino acid metabolism, nucleotide metabolism, tRNA synthesis, ribosomal protein, transporter and permease genes). We also found that Type III Secretion System effectors and transcription factors are part of the AlgU regulon (Table 1).

**Table 1.** Function groups of AlgU-induced genes in naïve leaves compared to King’s B media.

| Number of genes | Function |
| --- | --- |
| 36 | Hypothetical and unknown function |
| 26 | Transporter, permease, and lipoprotein |
| 23 | Amino acid, nucleotide, lipid metabolism, and energy production |
| 19 | T3SS related |
| 19 | Transcription regulator |
| 16 | tRNA |
| 14 | Protein translation and folding |
| 13 | Alginate synthesis |
| 10 | Ribosomal protein |
| 9 | Osmotic stress |
| 6 | T2SS structural |
| 6 | Oxidative response |
| 5 | Cell shape maintenance and division |
| 5 | DNA repair |
| 4 | Coronatine synthesis |
| 3 | Phage related |
| 2 | Iron-sulfur cluster related |
| 2 | Other secretion system |
| 2 | SOS response |
| 2 | Antibiotic resistance and toxin/antitoxin |
| 2 | Plasmid mobility gene |

For the AlgU downregulated *in planta* regulon, we identified genes involved in flagellar assembly, chemotaxis, signal transduction pathways, Type IV conjugation pilus assembly, DNA modification pathways, and macromolecule metabolism and energy production (including lipid metabolism, amino acid metabolism, nucleotide metabolism, protein translation and degradation, and transporter genes). We also identified GGDEF/EAL domain containing proteins and transcription factors among downregulated genes (Table 2).

**Table 2.** Function groups of AlgU down-regulated genes in naïve leaves.

| Number of genes | Function |
| --- | --- |
| 35 | Hypothetical and unknown function |
| 32 | Amino acid, nucleotide, and lipid metabolism, and energy production |
| 14 | Transporter, permease, and lipoprotein |
| 13 | Transcription regulator |
| 11 | Flagella related |
| 10 | Chemotaxis |
| 10 | Receptor and signal transduction |
| 8 | Conjugal transfer protein |
| 6 | Protein translation, folding, and protease |
| 5 | Phage or transposase |
| 3 | GGDEF domain containing protein |
| 3 | DNA modification |
| 3 | Sulfur metabolism or iron-sulfur cluster related |
| 1 | Antibiotic synthesis |

### Pre-induced PTI response intervenes against the AlgU *in planta* regulon

It was previously observed that the *Pto* genes affected by pre-induced PTI include those that are regulated by AlgU. To further identify the portion of the AlgU *in planta* regulon that is affected by PTI, we compared the AlgU *in planta* regulon to the list of genes that are affected by PTI (Lovelace et al. 2018). We found that 124 (55%) genes from the AlgU upregulated *in planta* regulon, and 91 (59%) genes from the AlgU downregulated *in planta* regulon are intervened against by the pre-induced PTI (Fig. 1B), which involves all the functional groups except iron-sulfur cluster related genes (Table 3, S3, and S4). The majority of the remaining AlgU *in planta* regulon genes remain the similar expression level under pre-induced PTI conditions compare to naïve conditions, namely 86 (38%) genes from the AlgU upregulated *in planta* regulon, 63 (41%) from the AlgU downregulated *in planta* regulon. In contrast, 14 (6%) genes in the AlgU upregulated *in planta* regulon are further upregulated under pre-induced PTI condition, and none of the AlgU-suppressed *in planta* regulon genes is further downregulated (Table S5). Taken together, these numbers suggest that the pre-induced PTI response and AlgU *in planta* regulon are in conflict.

**Table 3.** PTI intervenes almost all AlgU-regulated function groups.

| Percent of *in planta* regulon<sup>a</sup> | Number of genes | Function |
| --- | --- | --- |
| 61.54% | 16 | Transporter, permease, and lipoprotein |
| 100.00% | 13 | Alginate synthesis |
| 36.11% | 13 | Hypothetical and unknown function |
| 52.17% | 12 | Amino acid, nucleotide, lipid metabolism, and energy production |
| 57.89% | 11 | T3SS related |
| 78.57% | 11 | Protein translation and folding |
| 90.00% | 9 | Ribosomal protein |
| 47.37% | 9 | Transcription regulator |
| 88.89% | 8 | Osmotic stress |
| 100.00% | 4 | Coronatine synthesis |
| 50.00% | 3 | T2SS structural |
| 100.00% | 3 | Phage related |
| 40.00% | 2 | Cell shape maintenance and division |
| 12.50% | 2 | tRNA |
| 100.00% | 2 | Other secretion system |
| 40.00% | 2 | DNA repair |
| 33.33% | 2 | Oxidative response |
| 50.00% | 1 | Antibiotic resistance and toxin/antitoxin |
| 50.00% | 1 | Plasmid mobility gene |
| 0.00% | 0 | Iron-sulfur cluster related |
| 0.00% | 0 | SOS response |

AlgU down-regulated genes:
| Percent of *in planta* regulon | Number of genes | Function |
| --- | --- | --- |
| 53.13% | 17 | Amino acid, nucleotide, and lipid metabolism, and energy production |
| 45.71% | 16 | Hypothetical and unknown function |
| 90.91% | 10 | Flagella related |
| 71.43% | 10 | Transporter, permease, and lipoprotein |
| 76.92% | 10 | Transcription regulator |
| 90.00% | 9 | Chemotaxis |
| 70.00% | 7 | Receptor and signal transduction |
| 50.00% | 3 | Protein translation, folding, and protease |
| 66.67% | 2 | GGDEF domain containing protein |
| 40.00% | 2 | Phage or transposase |
| 66.67% | 2 | Sulfur metabolism or iron-sulfur cluster related |
| 100.00% | 1 | Antibiotic synthesis |
| 12.50% | 1 | Conjugal transfer protein |
| 33.33% | 1 | DNA modification |

a. Percent of genes in each function groups in Table 1 or 2 that are intervened by PTI.

### Stress responsive genes are AlgU induced and intervened against by PTI at 5 hpi

AlgU regulates the expression of osmotic and oxidative stress response genes in different bacteria (Wang et al. 2021). Alginate, a secreted polysaccharide, is also generally considered to shield bacteria from external stressors (Chang et al. 2007; Keith et al. 2003). We analyzed the relationship between AlgU and these stress tolerance related genes *in planta*. Glycine betaine transporter genes (*opuCABCD, cbcVWX*)(Chen and Beattie 2007; Chen et al. 2010) were dependent on AlgU for induction, and are PTI inhibited (Fig. 2A). However, for compatible solute synthesis genes, even though their induction is AlgU dependent, their expression was not inhibited by the pre-induced PTI at 5 hpi. The oxidative stress response genes (according to Yu 2014) *trx-2*(PSPTO_5243), *sodB* (PSPTO_4363), and a glutaredoxin domain protein (PSPTO_4161) are also part of the AlgU *in planta* regulon that are intervened against by PTI (Table S1 and S3). Alginate synthesis genes showed a similar trend to the glycine betaine transporter genes, they are induced by AlgU, and intervened against by pre-induced PTI (Fig. 2B). These results suggest that at 5 hpi, the bacterial cells activate genes related to osmotic stress response and alginate production with the help of AlgU, while pre-induced PTI can have a negative effect on the induction, which likely reduces the bacterial capability for tolerating stresses.

**Figure 2.**
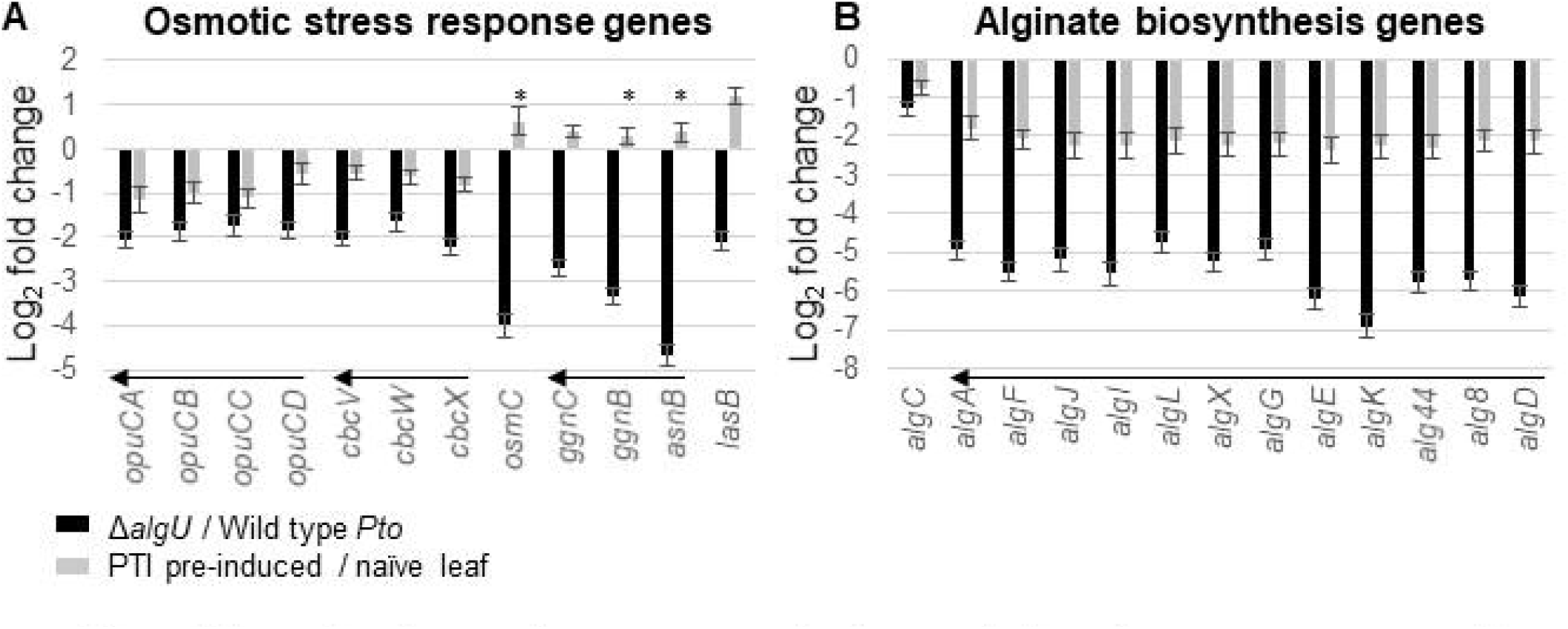
Expression changes of stress response related genes. **A**. Osmotic stress response genes. **B**. Alginate synthesis genes. * indicates genes with padj >0.05 calculated by DESeq2. All * in this graph are from grey bars. Arrows indicate genes within an operon.

### Multiple secretion system-associated genes are AlgU induced and intervened against by PTI at 5hpi

The Type III Secretion System (T3SS) is a needle-like structure that secrets unfolded effector proteins, and is required for *Pto* virulence (O’Malley and Anderson 2021; Lindgren et al. 1986). Type III effectors (T3Es) are delivered into the host cells and are critical for suppressing PTI responses in naïve plants. Genes encoding T3Es are distributed at more than ten locations in the *Pto* genome (Cunnac et al. 2011; Wei et al. 2015). Our data showed that AlgU enhances induction of 17 out of the 36 T3Es at 5 hpi, and these same genes are suppressed by pre-induced PTI (Fig. 3). However, T3SS structural genes are not significantly AlgU induced, even though they are suppressed by pre-induced PTI (Fig. S2).

**Figure 3.**
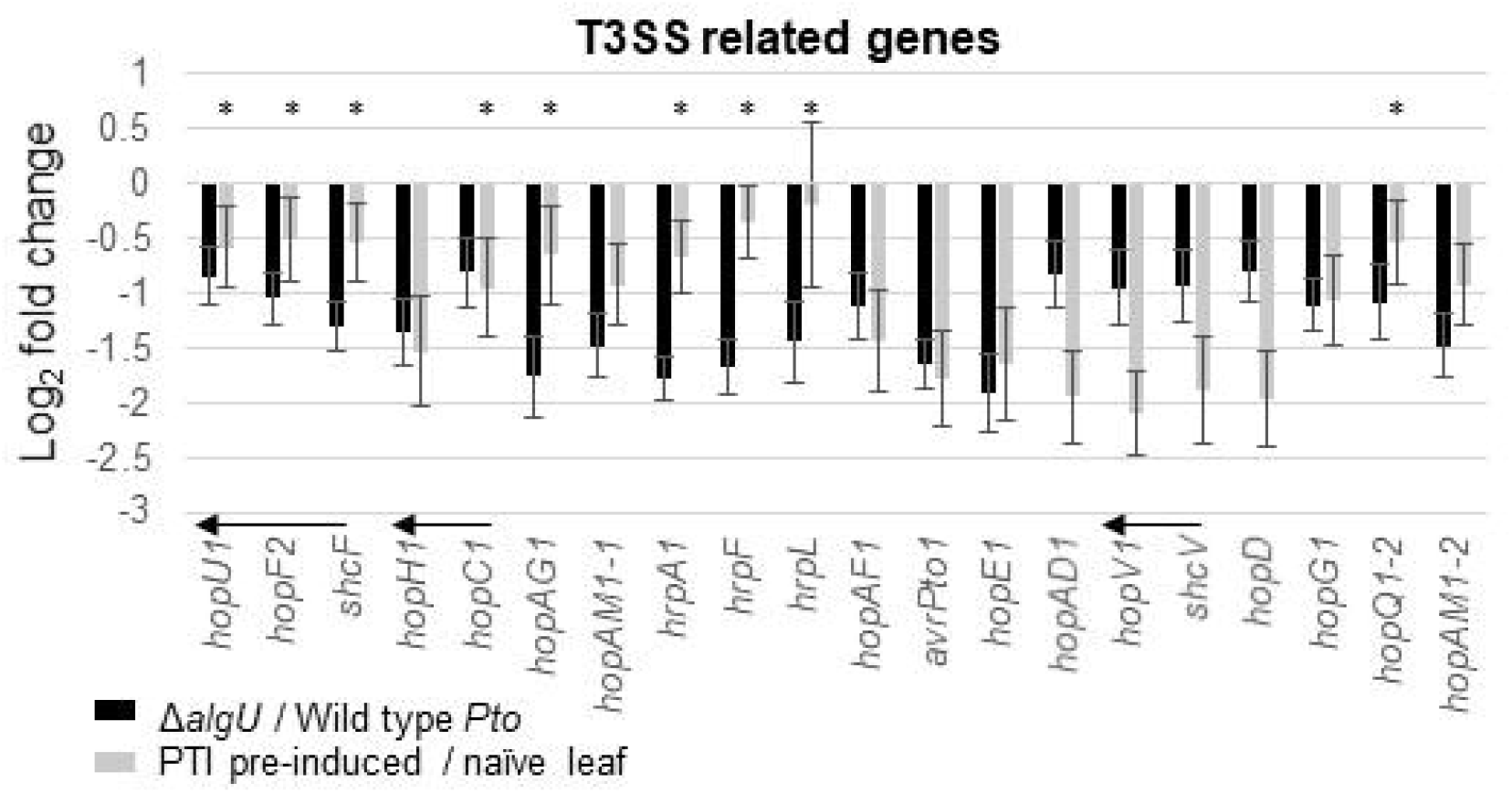
Expression changes of Type 3 Effectors and Type 3 Secretion System (T3SS) related genes that were identified as AlgU regulated in this study. * indicates genes with padj >0.05 calculated by DESeq2. All * in this graph are from grey bars. Arrows indicate genes within an operon.

Another secretion system that may also play a role in PTI response manipulation, the Type II Secretion System (T2SS), transports folded proteins from periplasm to the extracellular space. Our data showed that ten T2SS structural component genes are AlgU induced at 5 hpi, and *gspD, gspN*, and *gspM* are suppressed in the pre-induced PTI environment (Fig. 4). Interestingly, *gspDNM* are the three last genes within the cluster, and the first several genes in the same cluster are not significantly PTI-suppressed. One explanation is that these genes may be transcribed from separate promoters.

**Figure 4.**
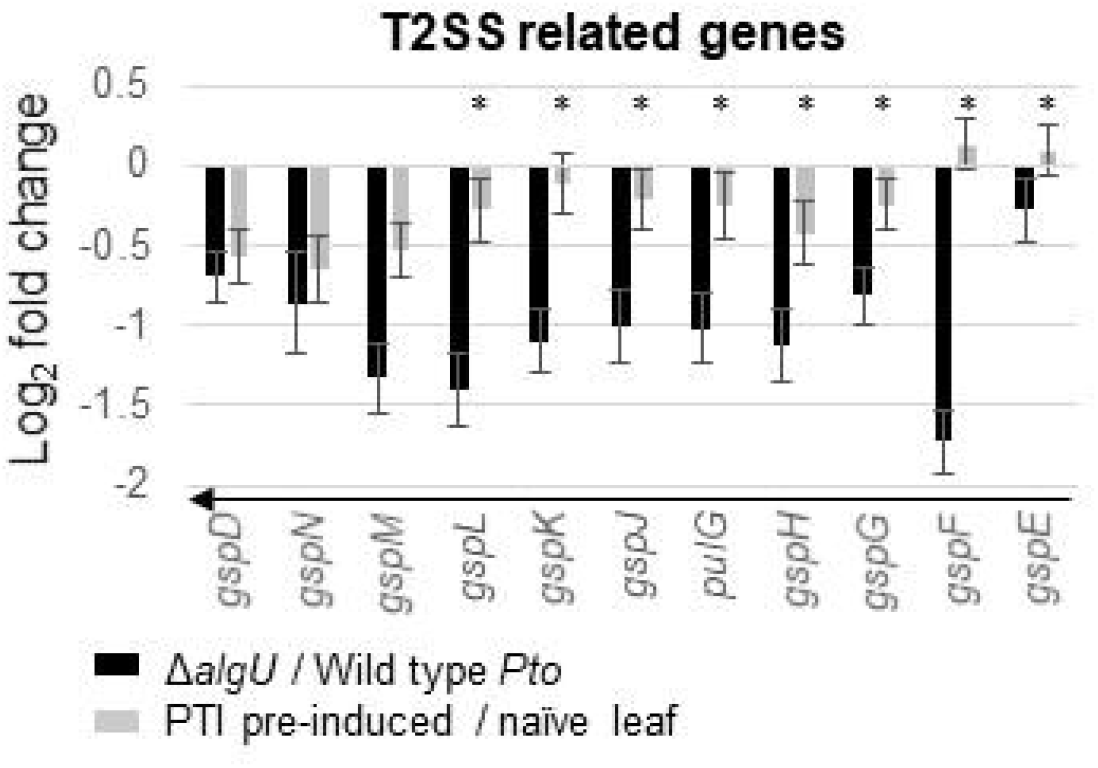
Expression changes of Type 2 Secretion System (T2SS) pathway genes. * indicates genes with padj >0.05 calculated by DESeq2. All * in this graph are from grey bars. Arrows indicate genes within an operon.

Coronatine is a plant hormone mimic and virulence factor produced by *Pto*. It was shown previously that AlgU may play a role in coronatine synthesis gene regulation (Ishiga et al. 2018). Our data showed that, at 5 hpi *in planta*, AlgU significantly upregulated the expression of only two coronatine-associated genes: *hopAQ1,* which may be co-transcribed with the regulator for coronafacic acid (CFA) and coronamic acid synthesis (CMA) called *corRS*, but is not known to directly relate to coronatine synthesis, and *cmaL*, which is encoded away from the primary biosynthetic CFA and CMA operons (Worley et al. 2013)(Fig. S3). Regardless of the limited involvement of AlgU in coronatine gene regulation, most of the coronatine biosynthesis genes are inhibited by pre-induced PTI.

### Flagellar motility-related genes are AlgU suppressed and intervened against by PTI at 5 hpi

In rich media, AlgU suppresses swimming motility by lowering the expression of several flagella-related genes including the *fliC* flagellin gene. Reduced flagellin expression can reduce FLS2-mediated responses in tobacco and tomato (Bao et al. 2020). We sought to answer if AlgU regulates motility related genes in *Arabidopsis*. The 60+ flagellar assembly and chemotaxis related genes in *Pto* are organized as a large single gene cluster on the chromosome. Studies in *P. aeruginosa*, which possesses a syntenous cluster, suggested that these genes can be organized into four classes based on their expression hierarchy (Dasgupta et al. 2003) (Fig. 5A). Our data showed that the presence of AlgU in the WT background reduced expression of most of the four classes of genes at 5 hpi, and PTI-exposure increased the expression of all these genes (Fig. 5B).

**Figure 5.**
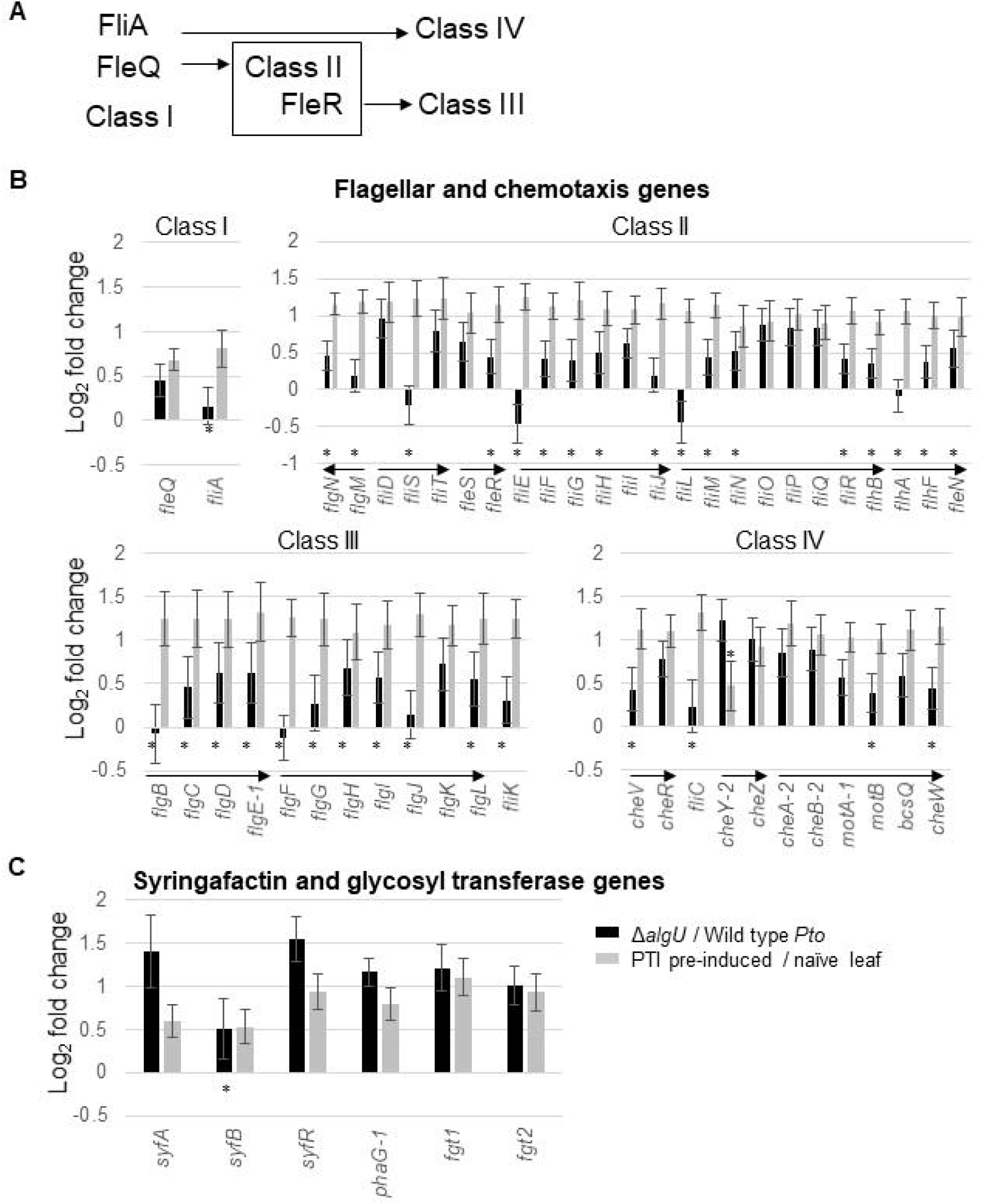
Expression changes of motility genes. **A**. Sketch showing the four classes belong to the motility gene regulatory hierarchy. **B**. Log2 fold change of the motility genes organized by classes. **C**. Genes related to swarming motility (syringafactin) and flagella glycosylation. * indicates genes with padj >0.05 calculated by DESeq2. * for black bars are placed under line of 0, * for grey bars are placed above line of 0. Arrows indicate genes within an operon.

Interestingly, several genes proposed to be involved in swarming motility showed the same trend as the four classes of flagellar genes (Fig. 5C). Syringafactin and 3-(3-hydroxyalkanoyloxy) alkanoic acid (HAA) are two surfactants, and their production is directly related to swarming motility (Nogales et al. 2015; Burch et al. 2012). The syringafactin production gene *syfA* and its transcription regulator *syrR(*also called *syfR)*, and the HAA production gene *rhlA* homolog *phaG-1* (PSPTO_3299) are all downregulated by AlgU and upregulated under pre-induced PTI conditions. In addition, the two genes *fgt1* and *fgt2* (PSPTO_1946/1947), which contribute to flagellar glycosylation and swarming motility (Taguchi et al. 2006), are similarly regulated. These results suggest that AlgU promotes the cells to enter a non-motile state at 5 hpi, while pre-induce PTI promotes the cells to keep expressing motility related genes.

### Multiple transcription factors have altered regulation *in planta* in the absence of AlgU, and are affected against by pre-induced PTI at 5 hpi

A previous study using ChIP-Seq has shown that many of the AlgU-regulated genes have no AlgU binding near their promoter region (Markel et al. 2016). One explanation is that AlgU affects the expression of these genes indirectly through other transcriptional regulators, either by direct induction, or through feedbacks. Our data showed that 19 genes with known or predicted transcriptional regulatory function, including *hrpL*, *amrZ*, and *fur* are induced when AlgU is present, and another 13 transcriptional regulators including *fleQ* and *syrR* are suppressed in the presence of AlgU at 5 hpi (Fig. 6). Pre-induced PTI significantly affected 9 of these induced genes and 10 of these suppressed genes. Interestingly, *amrZ* showed a strong AlgU dependence for induction, but it is induced under PTI condition rather than suppressed, dissimilar to the others.

**Figure 6.**
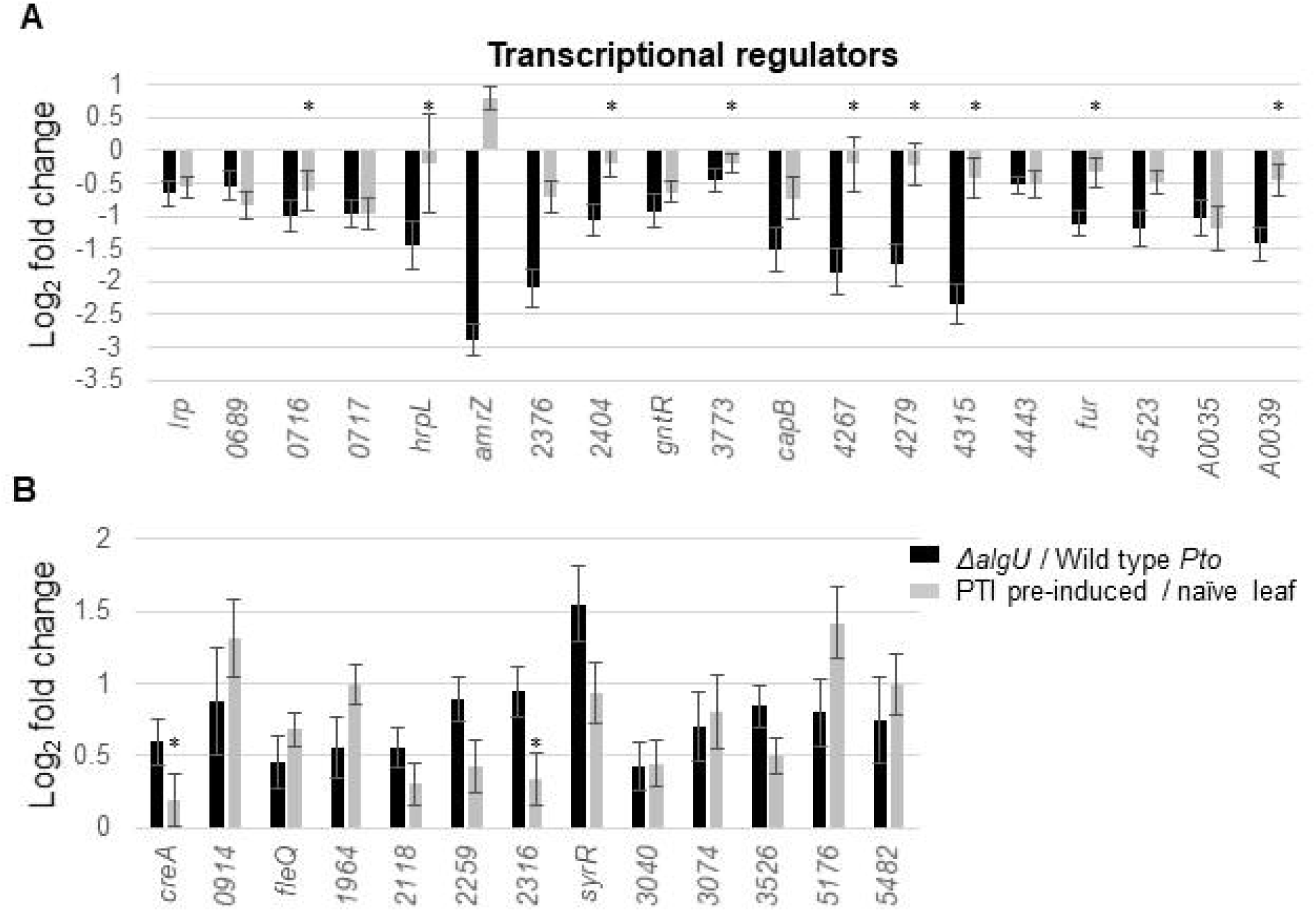
Expression changes of transcriptional regulators that are differentially-expressed in the absence of AlgU in naïve plant. **A**. Genes downregulated in *ΔalgU* background. **B**. Genes upregulated in *ΔalgU* background. * indicates genes with padj >0.05 calculated by DESeq2. All * in this graph are from grey bars.

## Discussion

In this study, we compared transcriptomes from *P. syringae* pv. *tomato* DC3000 WT and *ΔalgU* strains, either at 5 hpi in four-week-old *Arabidopsis thaliana* Col-0 with or without flg22 pre-treatment, or in KB media, to find AlgU-regulated genes specific to the apoplastic niche, which we termed “AlgU *in planta* regulon”. We then compared the transcriptome profile of these genes under pre-induced PTI and naïve conditions and determined that PTI affects more than half of AlgU-regulated genes, and that PTI affects almost all function groups regulated by AlgU. Our result suggests that AlgU and PTI-mediated regulation may be in conflict.

### AlgU *in planta* regulon is smaller from that was observed *in vitro*

The number of genes in the AlgU *in planta* regulon is smaller than the published AlgU *in vitro* regulon (Markel et al. 2016), which was generated using an AlgU over-producing strain in King’s B media. Our study looked at native expression, but not over-expression, which may account for the difference between the two results. The smaller size of the AlgU *in planta* has also been observed from a previous study, which looked at the AlgU regulon under different *in vitro* stressors or in association with plant hosts and showed that subsets of many transcription factor regulons are condition specific (Yu et al. 2014). It is possible that in the plant niche, a subset of the transcriptional factors in the AlgU regulon can incorporate condition specific signals. Another explanation for the environment-specific regulation is that AlgU regulon genes are co-regulated by other transcription factors. Previous studies and our data have shown that the AlgU regulon overlaps with other transcription regulators such as the sigma factors RpoS, RpoN, HrpL, (Markel et al. 2016; Yu et al. 2014), small RNA regulators RsmA2/A3 (Liu et al. 2021), the Fur iron homeostasis regulator (Butcher et al. 2011; Nobori et al. 2018), and the two-component system CvsSR (Fishman et al. 2018).

### Secretion systems

Consistent with the AlgU *in vitro* regulon, we have identified multiple T3Es as part of the AlgU *in planta* regulon. However, we could not find common regulatory or functional features among these T3Es to explain why only a specific subset of effectors are AlgU regulated. These T3E are from different clusters on the genome, each has a different identified function in the plant cell, and each interact with different plant proteins. Some T3Es targets both PTI and Effector Triggered Immunity (ETI) pathways (HopAD1 (Wei et al. 2015) and HopU1 (Nicaise et al. 2013; Fu et al. 2007)). Some T3Es are associated with cytoskeleton (HopG1 (Shimono et al. 2016; Block et al. 2010) and HopE1 (Guo et al. 2016)). Some T3Es are associated with plant membrane proteins (HopF2 (Zhou et al. 2014) and HopAF1 (Washington et al. 2016)). The only shared feature of these T3Es is that they suppress PTI. However, PTI suppression is a common feature of T3Es and many of them are not a part of the AlgU *in planta* regulon. The sigma factor HrpL, which regulates the expression of T3SS related genes, was identified as an AlgU *in planta* regulated gene, similarly to the observation from previous studies (Markel et al. 2016; Yu et al. 2014). It was previously shown that *hrpL* is downregulated by PTI compared to naïve plants at 1 hpi and 3 hpi during infection, but is not significantly differentially regulated at 5 hpi, (Lovelace et al. 2018).

Unlike the T3SS, the importance of T2SS in the host-pathogen relationship has been overlooked in *Pto,* which is a hemibiotrophic pathogen. In agreement with the previously published AlgU regulon in growth media (Markel et al. 2016), our data also showed that T2SS structural genes are upregulated by AlgU *in planta,* and these genes are PTI suppressed. T2SS is a part of the general secretion pathway (GSP) and transports folded proteins across the outer membrane. Even though it was shown that a significant reduction in disease development happened in both *Δ*g*spD* and *ΔgspE* background (Bronstein et al. 2005), most research regarding to virulence factor secretion in *Pto* focused on the T3SS. In contrast to *Pto*, T2SS is better understood in several other pathogens. In the necrotrophic pathogens *Pectobacterium atrosepticum* and *Dickeya dadantii,* T2SS is important for virulence, ROS tolerance (Charkowski et al. 2012; Liu et al. 2019), and promotes commensal bacterial growth via secretion of pectate lyases and other cell-wall-degrading enzymes (Yamazaki et al. 2011). Similarly, in the root commensal bacteria *Dyella japonica,* T2SS plays a role in PTI suppression (Teixeira et al. 2021). In the human pathogen *P. aeruginosa*, T2SS is involved in biofilm formation (Lewenza et al. 2017) and toxin secretion (Swietnicki et al. 2019). On the other hand, even though our data showed that T2SS is AlgU induced *in planta* and is PTI-suppressed, we do not know which proteins are T2SS substrates, because the substrate recognition signal is not universal and cannot be easily predicted (Naskar et al. 2021; Pineau et al. 2014). The two known substrates PlcA1 (PSPTO_3648) and PlcA2 (PSPTO_B0005) (Bronstein et al. 2005) are neither significantly induced by AlgU nor affected by pre-induced PTI (data not shown). Based on research from other bacteria, the substrates secreted by T2SS in *Pto* may include lipases, proteases, phosphatases, and more (Tilley et al. 2014; Urusova et al. 2019; Putker et al. 2013).

Interestingly, in contrast to a previous study which identified T6SS genes as a part of the *P. syringae* pv. *syringae* B728a AlgU regulon in the bean apoplast (Yu et al. 2014), our result did not identify any T6SS genes. There are two copies of T6SS genes, the first copy HSI-I coded by genes PSPTO_2538-2554, and the second copy HSI-II coded by PSPTO_5415-5438 (Sarris et al. 2010). Both copies include both structural genes and secreted factors. Our data did not identify any of these genes as part of the AlgU *in planta* regulon (data not shown). This is similar to the previous observation that *P. syringae* B728a and *Pto* have different T6SS regulation *in vitro* (Freeman 2013).

### Motility genes

It is interesting that we saw both swimming and swarming motility/surfactant related genes are suppressed by AlgU at 5 hpi, and are upregulated under the pre-induced PTI condition. This suggests that in addition to the previously proposed role for AlgU in immunity evasion, *Pto* may favor a nonmotile lifestyle in naïve plants. The enhanced expression of flagellar genes in the absence of AlgU has been shown to enhance immunity detection (Bao et al. 2020). It is interesting that exposure to PTI-conditions results in a similar pattern of enhanced flagellar gene expression by *Pto* which would presumably also enhance immune detection by the host. It is unclear whether this is a maladaptive response by the bacteria that favors the host or if PTI exposure signals are perceived by the bacteria as repellants resulting in a motility-driven avoidance response.

According to the flagellar gene regulatory hierarchy, the two transcription factors FleQ and FliA belong to class I. FleQ activates class II genes, which include the cytoplasmic structural genes and the regulator FleR. FleR then activates class III genes, which include the structural genes localized at the cell wall, outer membrane, and extracellular space. FliA activates class IV genes, which include flagellin, motor genes and chemotaxis genes (Dasgupta et al. 2003) (Fig. 5A). Interestingly, we observed two unexpected patterns from the RNA-seq data. First, even though PTI universally activated all these genes, the first gene in all FleQ and FleR induced operons showed distinctive AlgU inducibility in contrast to the other genes, as they are slightly induced while the following genes are downregulated by AlgU. Additionally, FleQ was hypothesized to suppresses the expression of the syringafactin regulator *syrR* (Nogales et al. 2015), but they both were suppressed by AlgU according to data from this study. These observations suggest that motility genes are not always co-regulated, and different regulatory mechanisms may exist. Second, *fliA* expression level was not significantly increased in the absence of AlgU, but most of FliA-regulated genes increased expression regardless. This may be because the change in expression level in the transcription regulator is not in proportion to the change in its regulated genes. Surprisingly, our RNA-seq data did not show a significant increase in the expression of *fliC* (Class IV) in the strain without *algU*, which is contradictory to the previously reported results based on RT-qPCR using tomato as the host at 6hpi (Bao et al. 2020). Instead, our data agrees with data from another group that also used *Arabidopsis* as the host plant (Ishiga et al. 2018). Whether the difference in *fliC* expression is due to differences in the host environment or other experimental factors remains an open question.

### Pre-induced PTI and AlgU in conflict

Overall, our data indicates that the patterns of induction or suppression for AlgU-regulated genes *in planta* are generally reversed during PTI exposure. At this point we are unable to conclude whether PTI-associated responses interfere with AlgU-mediated regulation directly or indirectly. The amount of free AlgU in the cell is post-translationally regulated by the anti-sigma factors MucA and MucB, which is regulated through the regulated intramembrane proteolysis (RIP) pathway. It was previously shown in *Arabidopsis* that two secreted plant proteases, SAP1 and SAP2, are induced during PTI and that these two proteases degraded the RIP pathway-associated protease MucD (Wang et al. 2019). However, in *P. syringae* and *P. aeruginosa,* a *ΔmucD* strain has a hyper-activated AlgU phenotype (Wang et al. 2019)(Damron and Yu 2011) while we have observed that PTI exposure has the opposite effect on AlgU-regulated genes. It is possible that AlgU is released via SAP1/SAP2 degradation of MucD and overactivation of the RIP cascade, but other PTI responses ultimately intervene against the AlgU regulon possibly through indirect overlapping regulators. The transcription factor AmrZ (PSPTO_1847), which was shown to have an AlgU binding site at its promoter region (Markel et al. 2016), is strongly dependent on AlgU for expression *in planta* (Fig. 6). AmrZ is one of the 14 AlgU upregulated genes that are also upregulated in PTI-activated plants compared to naïve plants (Table S5), supporting the possibility that AlgU is activated during PTI exposure. However, our data showed that these 14 genes make up less than 4% of the total AlgU *in planta* regulon, whether AlgU is activated and acted against, or if AlgU activity is directly suppressed by PTI remains a question. The specific cues within the apoplast that are perceived to induce the AlgU regulon are still unknown. Determining what signals bacteria perceive in the apoplast to drive AlgU activation and whether those signals are modified or masked during PTI will be critical to understanding how this conserved bacterial sigma factor interacts with this ancient form of plant immunity.

## Material and methods

### Plant material

*Arabidopsis thaliana* Col-0 seeds were sown in SunGrow Professional growing potting mix in 3.5-inch square pots, stratified for one day at 4°C in darkness, then moved to Conviron A1000 growth chamber with settings of 14 hour day, 23°C, 70 μmol light. After two weeks, the pots were thinned to 4 plants per pot. After four weeks, the pots were moved to a growth room with settings of 12 hour day, 23°C.

### RNA sample preparation and sequencing

The same procedure as described in a previous publication was used (Lovelace et al. 2018). At four to five weeks’ age, the four largest leaves of each plant were treated with flg22 or mock solution. Flg22 peptide (GenScript RP19986) stock solution was made by dissolving in DMSO to final concentration at 1mM. A 1000x dilution of the flg22 stock solution in water was syringe infiltrated to induce PTI, and a 1000x dilution of DMSO was used as the mock treatment for naïve conditions in this study. A 1mL blunt syringe was used to infiltrate the leaf from the abaxial side, through holes poked with a needle per half of the leaf. 24 leaves per timepoint per treatment were used. Each experiment was repeated 3 times and each produced a sequencing data set.

18-22 hours after the treatment, *Pto* was syringe-infiltrated into the treated leaves. To prepare the inoculum, *Pto* strains were first spread on King’s B Agar (King et al. 1954) supplemented with 20μg/mL rifampicin and grown to lawns overnight at room temperature. Cells from the lawn were harvested and resuspended in 0.25mM MgCl2 to an optical density 600nm of 0.8. The resuspension was used as the inoculum. After infiltration, excess liquid on the leaves were soaked up with paper towels. The bacteria strains used are listed in Table 4.

**Table 4.** Bacteria strains used.

| Stock # | Genotype | Plasmid | Reference |
| --- | --- | --- | --- |
| GS_00950 | *Pto* DC3000 Wild type | pJN105 | This study |
| GS_00585 | *Pto* DC3000 $\Delta algUmucAB$ | pJN105 | Markel et al. 2016 |

At 5 hpi, leaves were cut at the petiole-leaf blade junction. The leaves were then lined up in the middle of a sheet of parafilm. The parafilm was then folded so that the leaves are held between two layers of the parafilm with the cut side pointing at the folding line. Small openings were cut at the folding line before the assembly was rolled up from side to side and inserted into the barrel of 20-mL syringes. The syringes were put in 50mL centrifuge tubes and an RNA stabilizing buffer (Wit et al. 2012) was poured into the syringe. The tubes were then vacuumed at 96kPA for 2 min followed by a slow release. The vacuum procedure was repeated twice to infiltrate the leaves with the buffer. The excess buffer was discarded, then the tubes with syringes inside were centrifuged at 1,000x g for 10 minutes at 4°C to collect the apoplast wash fluid. The collected fluid was then passed through 0.20-μm Micropore Express Plus membrane filters (Millipore) to collect the bacteria. The filters were then placed in homogenization tubes and frozen by liquid nitrogen before storing at −80°C.

The filters were homogenized using Geno/Grinder (SPEX SamplePrep) for 1 min at 1,750 Hz on a liquid nitrogen chilled sample holder. Trizol (Thermo Fisher Scientific) was then added, and the Direct-Zol RNA Miniprep Plus Kit (Zymo Research) or Monarch RNA isolation kit (NEB) was used for RNA extraction. An additional TURBO DNase treatment (Ambion, Invitrogen/Thermo Fisher) was carried out to further reduce DNA content, followed by a cleanup step using Monarch RNA cleanup kit (NEB). Finally, the library was produced using the TruSeq Stranded Total RNA library prep kit (Illumina). RNA sample was then sent to Georgia Genomics and Bioinformatics Core for quantification, QC analysis and RNA sequencing. Single-end 75nt reads were sequenced using Nextseq 500 system (Illumina) in high output mode.

### Data analysis

The sequencing reads were first trimmed using Trimmomatic V0.36 and read counts were computed using EDGE-pro V1.3.1 and exported using edgeToDeseq.perl. Differentially-expressed genes were then identified using DESeq2 V1.28.1, with an adjusted P value below 0.05 and a log fold change over 0.58. Venn diagram was generated with InteractiVenn website (Heberle et al. 2015) and meta-chart website. Illumina sequencing data were deposited in the Gene Expression Omnibus under accession number GSE191032.

Protein function groups were manually curated by combining the outcome of Kyoto Encyclopedia of Genes and Genomes (KEGG), NCBI protein, Uniprot, pseudomonas.com, and literature mining. Protein cytoplasm or membrane localization was determined by checking pseudomonas.com and Uniprot.

### RT-qPCR

For real time quantification PCR analysis of selected genes, plants were treated in the same way as described above. At 5 hpi, three 0.4mm diameter leaf disks from inoculated leaves were taken and frozen by liquid nitrogen. Leaves were then crushed by Geno/Grinder in liquid nitrogen chilled sample holder. Then Trizol was added before RNA was extracted using Monarch RNA Cleanup Kit (NEB), skipping the gDNA cleaning column steps. After the extraction, the sample was treated with TURBO DNase. Monarch total RNA miniprep kit was then used to clean up the reaction. The RNA was then reverse-transcribed using QuantaBio qScript cDNA SuperMix. Luna qPCR Master Mix (NEB) was used for reaction setup, and StepOnePlus (Applied Biosystems) was used to carry out the PCR reaction. Reference genes (*hemD, Isc-1,* and 16s rRNA) were chosen from the list published previously (Smith et al. 2018), and showed minimal change in expression between conditions, with expression level close to genes of interest, based on the RNA-seq data. Raw data from StepOnePlus was then analyzed using LinRegPCR, and expression fold change was calculated using Microsoft Excel. Primers used are listed in table 5.

**Table 5.**
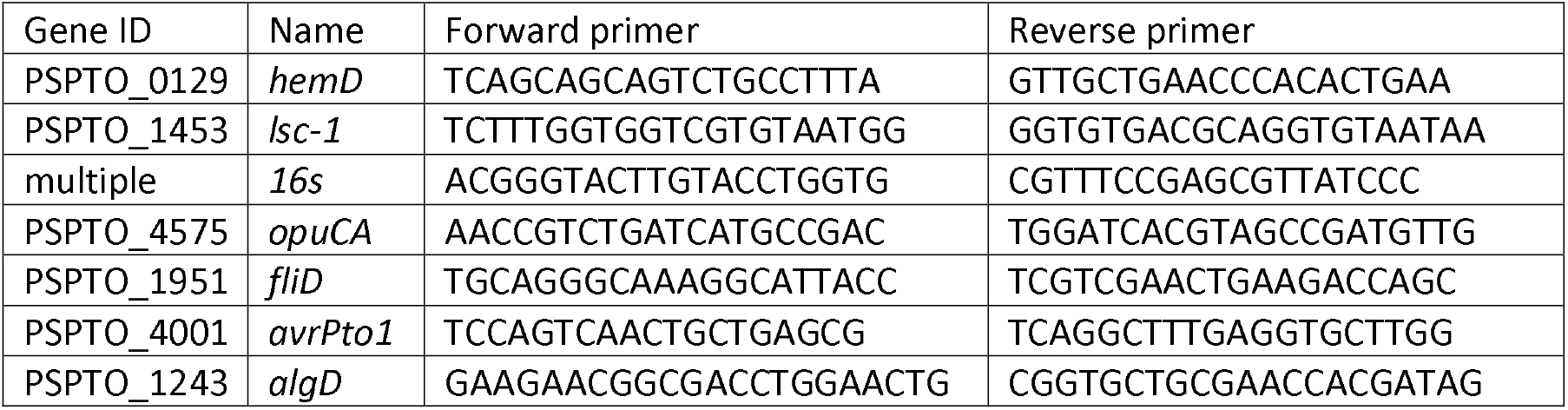
Primers used for RT-qPCR. *hemD*, *lsc-1*, and 16s primers were used in Smith et al. 2018 and were used as reference genes in this study. Primer sequences of *algD* is from Markel et al. 2016. Gene IDs are based on GenBank Pto genome NC_004578.1.

## Acknowledgements

We thank Dr. Bryan Swingle at Cornell University for providing strains and providing critical feedback on the drafting of this manuscript. We would also like to thank the members of the Kvitko lab and lab of Dr. Li Yang, and Dr. Mei Zhao at UGA for helpful discussions regarding the preparation of the manuscript.

