## Supplementary figures and images for "*In planta* transcriptomics reveals conflicts between pattern-triggered immunity and the AlgU sigma factor regulon"

### Figure S1

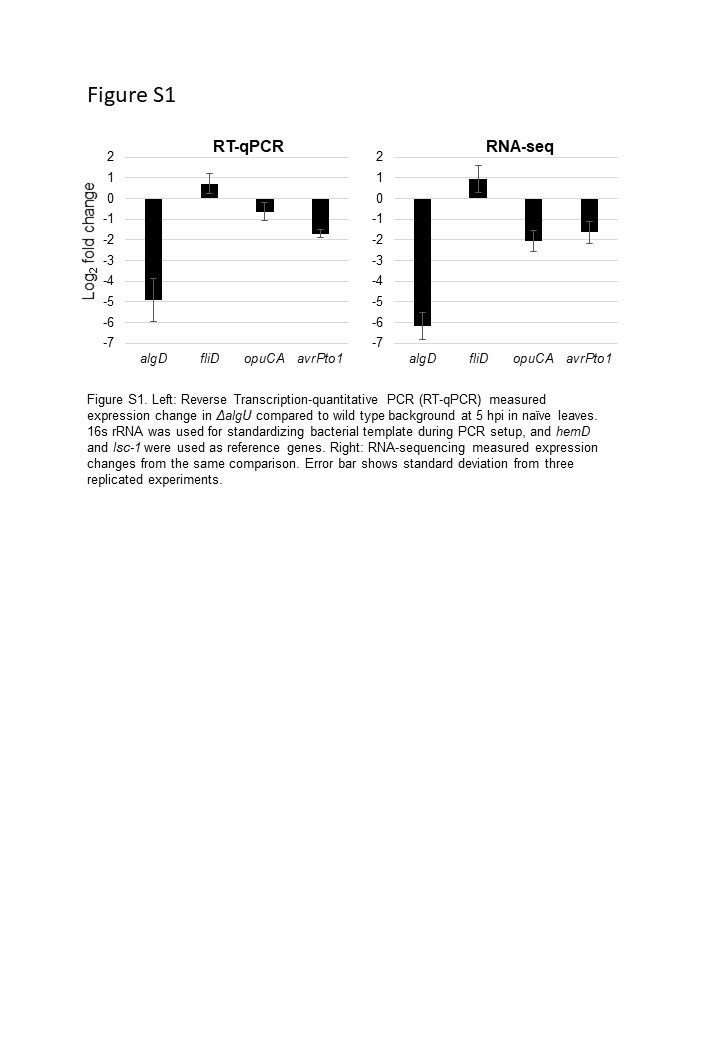

### Figure S2

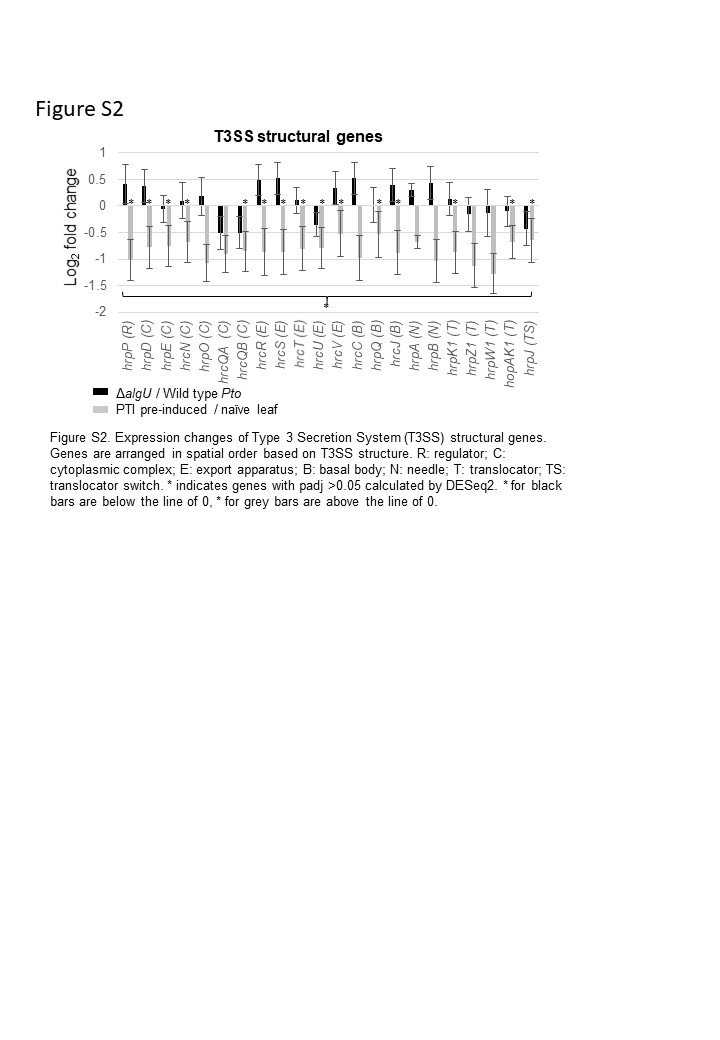

### Figure S3

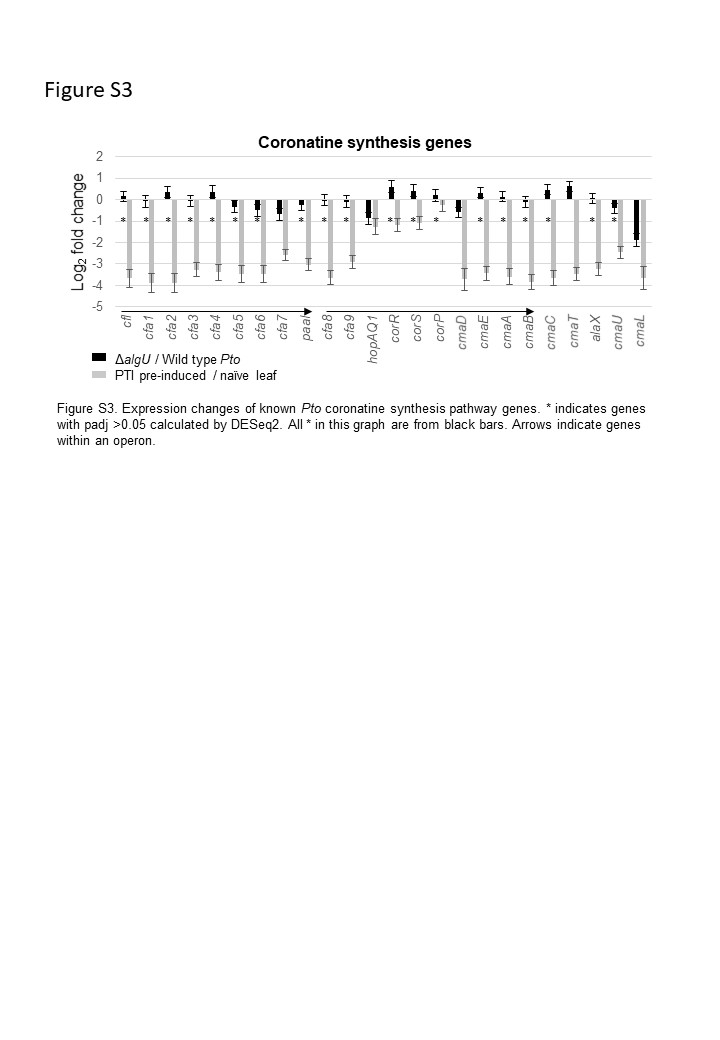
